# The pro-oncogenic adaptor CIN85 inhibits hypoxia-inducible factor prolyl hydroxylase-2

**DOI:** 10.1101/466946

**Authors:** Nina Kozlova, Daniela Mennerich, Anatoly Samoylenko, Elitsa Y. Dimova, Peppi Koivunen, Ekaterina Biterova, Kati Richter, Antti Hassinen, Sakari Kellokumpu, Aki Manninen, Ilkka Miinalainen, Virpi Glumoff, Lloyd Ruddock, Lyudmyla Drobot, Thomas Kietzmann

**Author notes:** Correspondence: Dr. Thomas Kietzmann, Faculty of Biochemistry and Molecular Medicine, University of Oulu, Aapistie 7C, 90220 Oulu, Finland.

## Abstract

The EGFR-adaptor protein CIN85 has been shown to promote breast cancer malignancy and hypoxia-inducible factor (HIF) stability. However, the mechanisms underlying cancer promotion remain ill-defined. Here, we show that CIN85 is a novel binding partner of the main HIF-prolyl hydroxylase PHD2, but not of PHD1 or PHD3. Mechanistically, the N-terminal SH3 domains of CIN85 interact with the proline-arginine rich region within the N-terminus of PHD2, thereby inhibiting PHD2 activity and HIF-degradation. This activity is essential in vivo, as specific loss of the CIN85-PHD2 interaction in CRISPR/Cas9 edited cells affected growth and migration properties as well as tumor growth in mice. Overall, we discovered a previously unrecognized tumor growth checkpoint that is regulated by CIN85-PHD2, and uncovered an essential survival function in tumor cells linking growth factor adaptors with hypoxia signaling.

## Introduction

Hypoxia-inducible factors (HIFs) are crucial in the adaption of tumor cells to the reduced availability of oxygen (1–3). From the HIFs known today, HIF-1α and HIF-2α appear to be essential promotors of malignant transformation, cell proliferation, invasion, and motility (4,5). Although several mechanisms contribute to the regulation of HIF-α expression (6,7), regulation of HIF stability by the family of HIF-prolyl hydroxylases (PHDs) is reported to be of major importance (8). Under normoxic conditions PHDs hydroxylate proline residues in HIF-1α and HIF-2α (9). This allows further recruitment of an E3-ubiquitin ligase complex containing the von-Hippel Lindau protein (VHL) (10), which results in the proteasomal degradation of HIF-1α and HIF-2α (11).

The pro-oncogenic adaptor protein CIN85 is a multi-modular scaffold protein, able of mediating various molecular interactions. Apart from being involved in downregulation of receptor tyrosine kinases such as EGFR (12,13), ErbB2/Her2 (14) and hepatocyte growth factor receptor (MET) (15). CIN85 also affects apoptosis (16,17), adhesion (18) and invasion (19). Earlier findings, including our own, showed that CIN85 promotes development of various cancers including breast cancer and displays highest levels in the most hypoxic areas of tumor tissues, which also usually display high HIF-α levels (20–22). In addition, we could previously show that CIN85 appears to induce HIF-1α stability via a so far unknown mechanism (23).

Thus, we hypothesized that CIN85 may affect HIF-α stability by affecting the HIF-PHDs. Accordingly, we examined the potential involvement of PHDs in CIN85-mediated HIF-α stability in a more mechanistic manner. Our current study clearly shows that CIN85 is a novel binding partner of the main HIF-hydroxylase PHD2. We further defined that the three SH3 domains of CIN85 and the proline-arginine rich area within the amino acids 77-100 in the N-terminus of PHD2 are of major importance for the CIN85-PHD2 complex formation. Additionally, we utilized a CRISPR/Cas9-mediated *EGLN1* (PHD2) gene editing approach to unravel the impact of the CIN85-PHD2 interaction on HIF-α stability and cellular behavior in triple negative breast cancer cells. The abrogation of PHD2-CIN85 complex formation in the CRISPR/Cas9-edited cells resulted in higher PHD2 activity and subsequently lower HIF-1α and HIF-2α levels, as well as in a less malignant cellular phenotype. Together, we show that CIN85 acts as a novel binding partner of PHD2, which can prevent prolyl-hydroxylation and degradation of HIF-α subunits, thereby promoting HIF-α stability and breast cancer malignancy.

## Results

### CIN85 and PHD2 undergo a direct interaction

In a previous study, we showed that CIN85 is able to induce HIF-1α stabilization and expression of its target gene plasminogen activator inhibitor-1 (PAI-1) (23). Therefore, we hypothesized that CIN85 contributes to HIF-1α stabilization by interfering with the function of PHDs via a direct interaction and explored this in more detail. Since PHD2 is the major variant regulating HIF-1α levels (24), we first performed co-immunoprecipitation studies in the three triple-negative breast cancer cell lines MDA-MB-231, BT-549, and Hs 578T. The results clearly show that endogenous CIN85 and PHD2 interact in all three cell lines (Figure 1A). Next, we investigated whether CIN85 is also able to interact with PHD1 and PHD3. In order to determine this, we performed co-immunoprecipitation studies in HEK-293 cells transiently overexpressing V5-tagged PHD1, PHD2, and PHD3 together with Myc-tagged CIN85. The co-immunoprecipitation assays revealed that only PHD2 binds to CIN85; no interaction of CIN85 with PHD1 or PHD3 could be detected (Figure 1B). Thus, these data indicate that CIN85 may interfere with HIF-α degradation only via PHD2 and not via PHD1 or PHD3.

**Figure 1.**
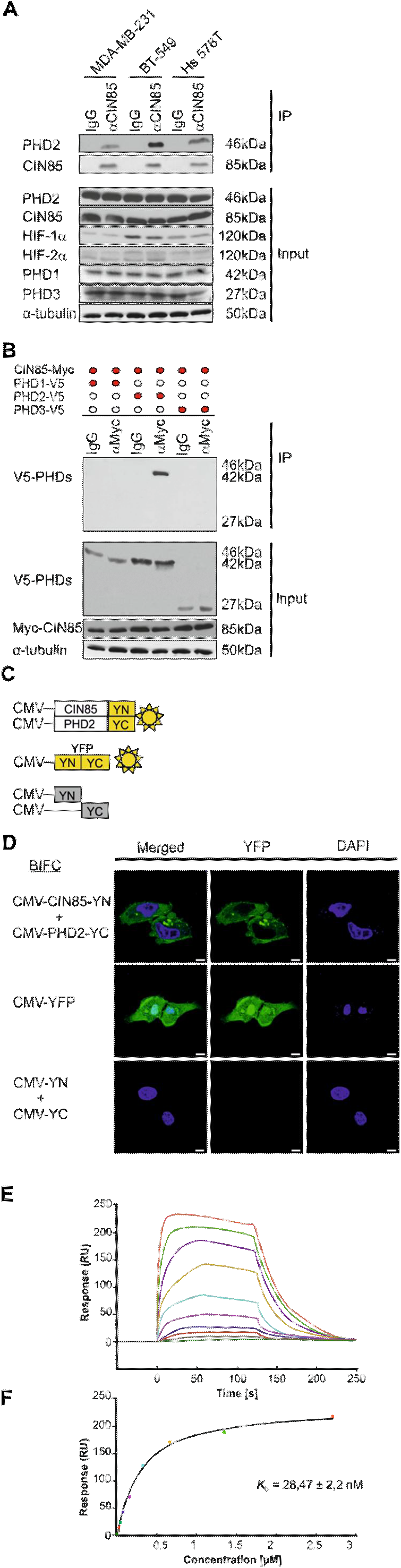
CIN85 interacts with PHD2. (A) Endogenous PHD2 was immunoprecipitated (IP) with CIN85 antibody from MDA-MB-231, BT-549, and Hs 578T cells. Blots from the input were probed with PHD1, −2, −3, CIN85, HIF-1α, and HIF-2α antibody. (B) Western blot (WB) analysis of immunoprecipitates (IP) and WCE from HEK-293 cells, expressing V5-tagged PHD1, PHD2, or PHD3 and Myc-tagged CIN85. Blots from IPs were probed with V5-tag antibody, the input was probed with V5-tag, Myc-tag and α-tubulin antibodies. (C) Schematic presentation of BiFC assay constructs. The constructs CMV-CIN85-YN and CMV-PHD2-YC allow expression of CIN85 and PHD2 as fusion proteins with the N-terminal or the C-terminal non-fluorescent parts of YFP (-YN and -YC) under the control of the CMV promoter, respectively. Note that CMV-YN and CMV-YC protein parts are non-fluorescent and non-interacting; however, interacting proteins such as CIN85 and PHD2 are able to reconstitute fluorescent YFP. (D) Visualization of the BiFC signal by confocal microscopy in BT-549 cells expressing CMV-CIN85-YN+CMV-PHD2-YC. No signal was detected upon expression of CMV-YN+CMV-YC. Scale bar 20 μm. (E) The surface plasmon resonance sensorgram of CIN85 binding to immobilized PHD2. Binding was assessed upon injection of a concentration series of CIN85 over immobilized PHD2. The CIN85 concentrations were 0, 10, 21, 42, 84, 168, 337, 675, 1350, 2700 nM (from bottom to top). (F) The fitted curve for different concentrations of CIN85 binding to immobilized PHD2 using the ‘Affinity’ model in the Biacore T200 evaluation software.

We next localized the PHD2-CIN85 complex in living cells by performing a bimolecular fluorescence complementation assay (BiFC). The BiFC assay is based on the formation of a fluorescent complex when two proteins, fused to the non-interacting and non-fluorescent parts of, in our case, the N-terminal and C-terminal halves of yellow-fluorescent protein (YFP), interact with each other (25,26). Upon that interaction, the YFP fluorescent complex is reconstituted and can be visualized. For this purpose, coding sequences of CIN85 and PHD2 were subcloned into pcDNA3-CMV-YN and pcDNA3-CMV-YC constructs, allowing the expression of PHD2 and CIN85 fused to the N-terminal or the C-terminal non-fluorescent parts of YFP, respectively. A description of the constructs used in the BiFC assay is schematically presented in Figure 1C. BT-549 cells were transfected with the CMV-CIN85-YN and CMV-PHD2-YC constructs. Transfection with a combination of unfused CMV-YN + CMV-YC constructs served as a negative control while transfection with a construct, encoding full length YFP served as a positive control. A punctated PHD2-CIN85 BiFC signal indicating the interaction of PHD2 with CIN85 could be visualized throughout the cytoplasm of the transfected cells (Figure 1D). Thus, these data indicate that PHD2 interacts with CIN85 in a non-nuclear compartment of the cells.

In order to understand whether the interaction between PHD2 and CIN85 is direct, we performed surface plasmon resonance (SPR) experiments. To this end, PHD2 was covalently immobilized to a SPR sensor chip, and a concentration series of recombinant CIN85 was injected over PHD2 and binding was assessed by SPR detection. The measurements revealed that full-length CIN85 bound to immobilized PHD2 with an apparent KD of 28,47 ± 2,2 nM (Figure 1E, F). Together, the data show that PHD2 and CIN85 interact in a direct manner.

### The three N-terminal SH3 domains of CIN85 bind to the N-terminus of PHD2

Next, we investigated which parts of each protein are involved in the interaction. CIN85 is a multi-domain adaptor protein consisting of 3 SH3 (SRC homology 3) domains (A, B and C), a proline rich domain (Pro) and a serine rich sequence (Ser), followed by a coiled-coil (CC) domain. PHD2 harbors a flexible N-terminus with the MYND (Myeloid, Nervy, and DEAF-1) Zn-finger domain and a catalytic domain on the C-terminus (Figure 2A).

**Figure 2.**
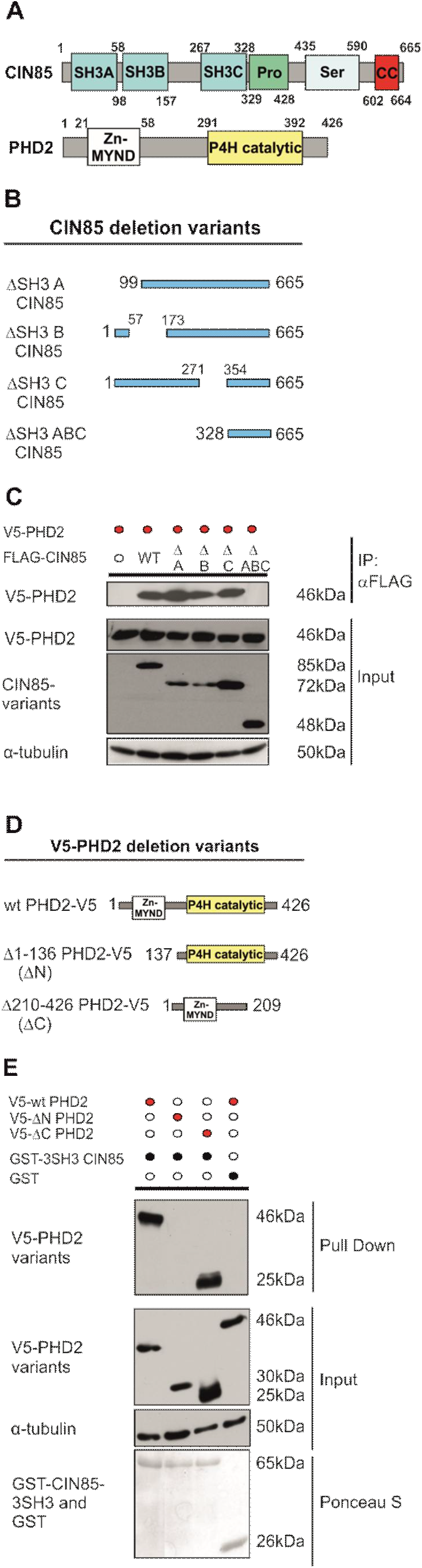
CIN85 interacts with the N-terminus of PHD2 via 3 SH3 domains. (A) Schematic presentation of the CIN85 and PHD2 structure. CIN85 consists of 3 N-terminal SH3 domains (A, B and C), a proline rich domain (Pro), and a serine rich sequence (Ser), followed by a coiled-coil domain (CC). The N-terminus of PHD2 possesses a MYND-Zn-finger domain, while the catalytic domain of PHD2 is located in the C-terminus. (B) Schematic presentation of FLAG-tagged CIN85 variants with a deletion of SH3A, SH3B, SH3C or all three SH3 domains used in the IP studies. (C) WB analysis of IPs and WCEs from HEK-293 cells, expressing V5-tagged PHD2 variants and FLAG-tagged-CIN85 variants. Blots from IPs were probed with V5-tag antibody, extracts were probed with V5-tag, FLAG-tag, and α-tubulin antibodies. (D) Schematic presentation of the V5-tagged PHD2 variants used in GST pull down (PD) studies: wt PHD2, PHD2 lacking amino acids 1-136 from the N-terminus (ΔN), and PHD2 variant lacking amino acids 210-426 from the C-terminus (ΔC). (E) WB analysis of PDs using recombinant GST or GST-fusion 3SH3 CIN85 protein in extracts from HEK-293 cells, expressing V5-tagged PHD2 variants (wt, ΔN, or ΔC). Blots from PDs were probed with V5-tag antibody; WCEs were probed with V5-tag and α-tubulin antibodies. Ponceau S stain of blots from PDs was used as a loading control.

First, we performed GST-pull down experiments with recombinant GST-SH3A, GST-SH3B, GST-SH3C, GST-Pro, GST-Ser, and GST-CC fusion proteins and lysates of HEK-293 cells transiently expressing V5-PHD2 (Supplementary Figure 1A). As a result, we were able to identify that the SH3 domains from CIN85 are responsible for the interaction with PHD2 (Supplementary Figure 1B), while no binding was observed between PHD2 and other domains of CIN85.

We next tested whether the CIN85-PHD2 interaction will be affected by the lack of a particular SH3 domain in CIN85. To do that co-immunoprecipitations with FLAG-tagged CIN85 lacking either SH3A, SH3B, SH3C or all three SH3 domains together with V5-tagged PHD2 were performed in HEK-293 cells (Figure 2B). Interestingly, we were able to observe that the CIN85-PHD2 interaction was present when CIN85 lacked either the SH3A, SH3B or SH3C domain, but the absence of all three SH3 domains abolished the interaction (Figure 2C). Collectively, these results indicate that the interaction between CIN85 and PHD2 depends on the presence of three N-terminal SH3 domains in CIN85.

In order to investigate which part of PHD2 participates in the binding to CIN85, we used, in addition to V5-tagged wild-type PHD2, two constructs that allow the expression of either a V5-tagged N-terminal part of PHD2 (V5-ΔC PHD2) or a V5-tagged C-terminal part (V5-ΔN PHD2) (Figure 2D). Additionally, since all three SH3 domains from CIN85 participate in the complex formation, we used a recombinant GST-3SH3-CIN85 protein in pull down studies as bait for the PHD2 deletion variants (Figure 2E). The results of the pull down assays show that GST-3SH3-CIN85 was able to interact with wt PHD2 and the PHD2 variant lacking the catalytic C-terminus (V5-ΔC PHD2). By contrast, no interaction was observed with a PHD2 lacking the first 136 amino acids of the N-terminus (V5-ΔN PHD2) (Figure 2E). Together, these data show that the interaction between PHD2 and CIN85 occurs between the three SH3 domains of CIN85 and the amino acids 1-136 within the N-terminus of PHD2.

### CIN85 binds a proline-arginine-rich sequence in the N-terminus of PHD2

In order to further delineate the CIN85-binding PHD2 region, we generated a series of PHD2 coding constructs lacking various amino acids from the N-terminus of PHD2 (Figure 3A) and GST-fusion proteins harboring peptides of different length from the PHD2 N-terminus (Supplementary figure 2A,C). The latter were used for pull-downs with lysates of HEK-293 cells transiently expressing full length Myc-CIN85 (Supplementary Figure 2A,2C). As a result, GST-1-40-PHD2 and GST-99-136-PHD2 were not able to pull down CIN85, indicating that amino acids 1–40 and 99–136 are not critical for the PHD2-CIN85 interaction (Supplementary Figure 2B). Other GST-PHD2 fragments used in the study were able to bind CIN85, although the binding was most evident when the amino acids 59-98 were present within the PHD2 fragment. Further pull down studies with shorter GST-PHD2 variants (Supplementary Figure 2C) revealed that GST-81-98-PHD2 showed the strongest binding to CIN85, while other GST-proteins were not able to pull down CIN85 to a similar level (Supplementary Figure 2D). Altogether the results of the pull down studies indicated that the PHD2 region between amino acids 58-98 is necessary for the binding to CIN85, while amino acids 81-98 have the strongest impact on the interaction with CIN85. Indeed, we observed similar results when using recombinant GST-3SH3-CIN85 as bait in the lysates of HEK-293 cells transiently expressing V5-tagged PHD2 variants lacking amino acids from the N-terminus (Figure 3A,B). While the PHD2 variants Δ1-26, Δ10-58, Δ99-136, and Δ210-426 were able to bind CIN85 like wt PHD2, the PHD2 variant lacking amino acids 52-98 showed an almost complete loss of binding like the PHD2 variant lacking the whole N-terminus (Figure 3A,B, Supplementary Figure 2E).

**Figure 3.**
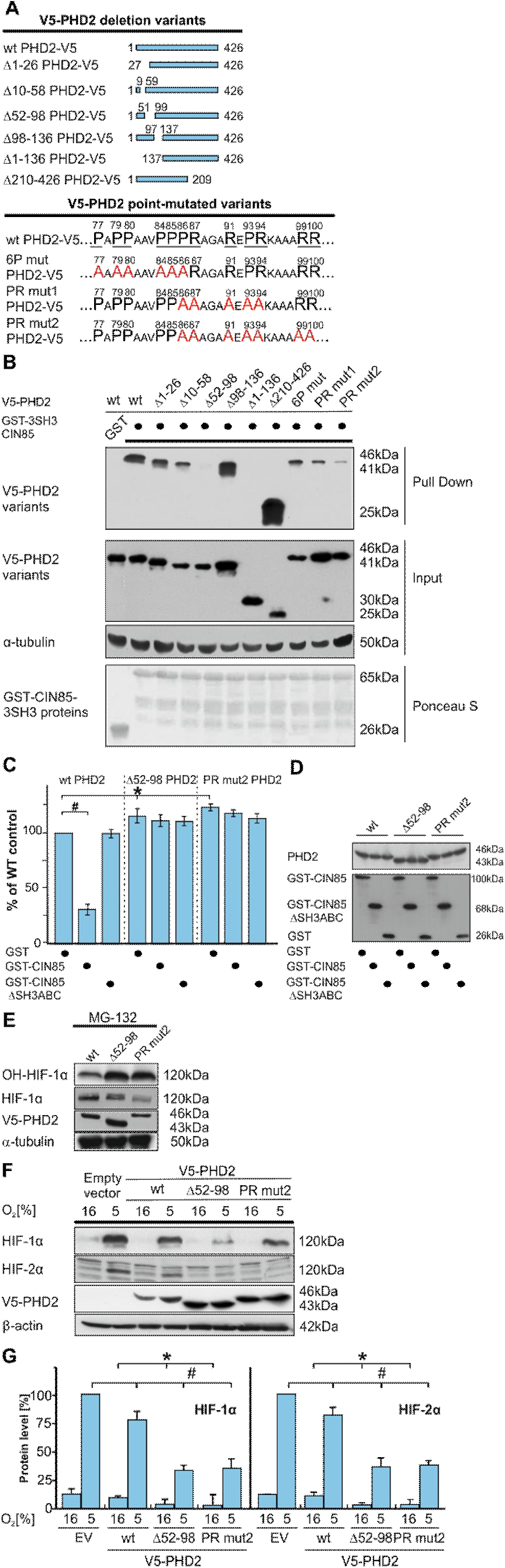
The PHD2 variants lacking the CIN85 binding pattern are of higher catalytic activity. (A) Schematic presentation of the V5-tagged PHD2 deletion variants and V5-tagged point-mutated PHD2 variants used in the PDs: wt PHD2, 1-26 (Δ1-26 PHD2), 10-58 (Δ10-58 PHD2), 52-98 (Δ52-98 PHD2), 99-136 (Δ99-136 PHD2), 1-136 (Δ1-136 PHD2), 209-426 (Δ210-426 PHD2), PHD2 P77A, P78A, P80A, P84A, P85A, P86A (6P), PHD2 P86A, R87A, R91 A, P93 A, R94A (PR mut1), and PHD2 P86 A, P87A, R91 A, P93 A, P94A, R99A, R100A (PR mut2). (B) WB analysis of PDs using recombinant GST-3SH3 CIN85 fusion proteins from HEK-293 cells expressing V5-tagged PHD2 variants. Blots from PDs were probed with V5-tag antibody; the input was probed with V5-tag and α-tubulin antibodies. Ponceau S stain of PDs was used as a loading control. (C) Results of the *in vitro* hydroxylation assay using wt PHD2, Δ10-58 PHD2, and PR mut2 PHD2 as catalysts and a [3H]proline-labeled HIF-1α peptide as a substrate in the presence of recombinant GST-CIN85 and GST-CIN85ΔSH3ABC. Recombinant GST was used as a control. The amount of 4-hydroxy[3H]proline formed was analyzed by a radiochemical assay (cf Materials and Methods) and used as a read out of the PHD2 activity. Wt PHD2+GST was set to 100%. Data are mean ±SD (n=3), * significant difference between Δ10-58 PHD2, and PR mut2 vs. wtPHD2. # significant difference between wtPHD2 and wtPHD2+CIN85. (D) Autoradiography of the [^35^S]Met labeled wt PHD2, Δ10-58 PHD2, and PR mut2 PHD2 used in the hydroxylation assay. (E) Hydroxy-HIF-1α levels in MG 132 treated cells expressing wt PHD2, Δ10-58 PHD2, and PR mut2 PHD2. (F) WB analysis of HEK-293 cells transiently expressing V5-tagged wt PHD2, Δ10-58 PHD2, and PR mut2 PHD2 cultured under normoxia (16%) or hypoxia (5% O_2_) for 6 h. Extracts were probed with HIF-1α, HIF-2α, V5, and α-tubulin antibodies. (G) Quantification of HIF-1α levels in HEK-293 cells transiently expressing empty control vector (EV), V5-tagged wt PHD2, Δ10-58 PHD2, and PR mut2 PHD2 cultured under normoxia or hypoxia. HIF-1α and HIF-2α levels in EV expressing cells cultured under hypoxia were set to 100%. Data are mean ±SD (n=4), * significant difference between Δ10-58 PHD2, and PR mut2 PHD2 expressing cells vs. wtPHD2 (normoxia). # significant difference between HIF-1α levels in wtPHD2, Δ10-58 PHD2, and PR mut2 PHD2 expressing cells vs. EV expressing cells (hypoxia).

The SH3 domains of CIN85 are reported to recognize atypical proline-arginine sequences (Kurakin, Wu et al. 2003) with each of the domains tolerating different amino acid adjacent to the arginine residue. To further unravel the nature of the CIN85-PHD2 binding we aimed to identify the proline and/or arginine residues within the region between amino acids 59-100 of PHD2 that are critical for the binding to CIN85. In order to verify their involvement in the binding, we created several V5-tagged PHD2 constructs where the respective proline or/and arginine residues were mutated to alanines (referred to 6P mut, PR mut1, and PR mut2) (Figure 3A). Pull down studies with these mutants and GST-3SH3-CIN85 revealed that these point-mutated PHD2 variants lost almost all their binding to CIN85 when compared to wt PHD2 (Fig. 3B). With the 6P mut variant where prolines 77, 78, 80, 84, 85, 86 were substituted with alanines binding was decreased by about 60%. When PR mut1 (P86, R87, R91, P93, R94 to alanine) was employed binding was reduced by about 80%. Lastly, the employment of PR mut2 (P86, P87, R91, P93, P94, R99, and R100 to alanine) reduced the binding between PHD2 and CIN85 by about 90% (Supplementary Figure 2E). Taken together, the results clearly indicate that the proline and arginine residues located within the area of amino acids 77–100 in the N-terminus of PHD2 are critical for the interaction with the SH3 domains of CIN85.

### PHD2 variants lacking CIN85 binding sequence show higher catalytic activity

Since our previous findings indicated that CIN85 stabilizes HIF-1α levels via interference with PHD-dependent HIF-1α protein degradation, we investigated whether the ability of PHD2 to interact with CIN85 has an impact on the catalytic activity of PHD2. The results of an *in vitro* HIF-1α peptide hydroxylation assay (cf Materials and methods) with recombinant PHD2 as catalyst revealed that addition of recombinant full-length CIN85 reduces the catalytic activity of PHD2 by about 70%. By contrast, a CIN85 variant lacking the three SH3 domains involved in the interaction with PHD2 did no longer reduce the PHD2 catalytic activity. In addition, when we used the PHD2 variants with no or the weakest binding to CIN85 (Δ52-98 PHD2; and PR mut2 PHD2, respectively) in the HIF-1α peptide hydroxylation assay we observed that both PHD2 variants showed a higher activity (about 15% and 20%, respectively) when compared to wt PHD2. Further, the activity of the PHD2 mutants was not affected by the presence of CIN85 (Figure 3C, D). In line with the results from the hydroxylation assay, overexpression of Δ52-98 and PR mut2 PHD2 increased the levels of hydroxy-HIF in MG123 treated cells (Figure 3E). Further, overexpression of Δ52-98 and PR mut2 PHD2 reduced HIF-1α levels under both normoxia and hypoxia, compared to wt PHD2 (Figure 3F, G). Together, these results suggest that lack of CIN85 binding promotes PHD2 activity, and subsequently lowers HIFα levels.

### Generation of MDA-MB-231 cells lacking the CIN85-PHD2 interaction

Next, we investigated the impact of the CIN85-PHD2 interaction *in vivo*. To address this, we used the CRISPR/Cas9 approach in triple negative MDA-MB-231 breast cancer cells in order to edit the *EGLN1* (PHD2) gene in a manner that it encodes a PHD2 lacking the amino acids required for binding to CIN85. By introducing two double strand breaks in *EGLN1* exon 1, this approach allowed deletion of 87 nucleotides (cf Materials and methods) (Supplementary Figure 3A).

The results of genotyping indicated the presence of various gene editing patterns after the introduction of Cas9 together with EGLN1-sgRNA-219 and EGLN1-sgRNA-292 into MDA-MB-231 cells (Supplementary Figure 3B). However, the respective PCR products from two clones (E10 and E12) appeared as a single band of the correct size on native PAGE. Further, we extracted total RNA from E10 and E12 cells and performed RT-PCR using the same set of genotyping primers. The PCR products from both clones appeared to be of shorter size compared to the PCR product from scrambled control cells (S) (Supplementary Figure 3C). Afterwards, the PCR products from the gDNA and cDNA of both clones were sequenced. As a result of sequencing, we observed only one type of template from both E10 and E12 MDA-MB-231 cells, which harbored a homozygous deletion of 87 bp and encodes a PHD2 variant lacking amino acids 75-103 (Supplementary Figure 3D). Next, by performing a Western Blot analysis, we were able to confirm that the deletion of 87 bp in exon 1 of *EGLN1* allowed the expression of a smaller Δ75-103 PHD2 when compared to the wt PHD2 in control cells (S) (Supplementary Figure 3E).

### Lack of amino acids 75-103 of PHD2 alters cell morphology and reduces HIF-α levels

As a next step, we performed immunoprecipitations and GST-pull downs using CIN85 antibodies and GST-3SH3-CIN85 and verified that the PHD2-CIN85 interaction in the E10 and E12 MDA-MB-231 cells was lost (Figure 4A, B).

**Figure 4.**
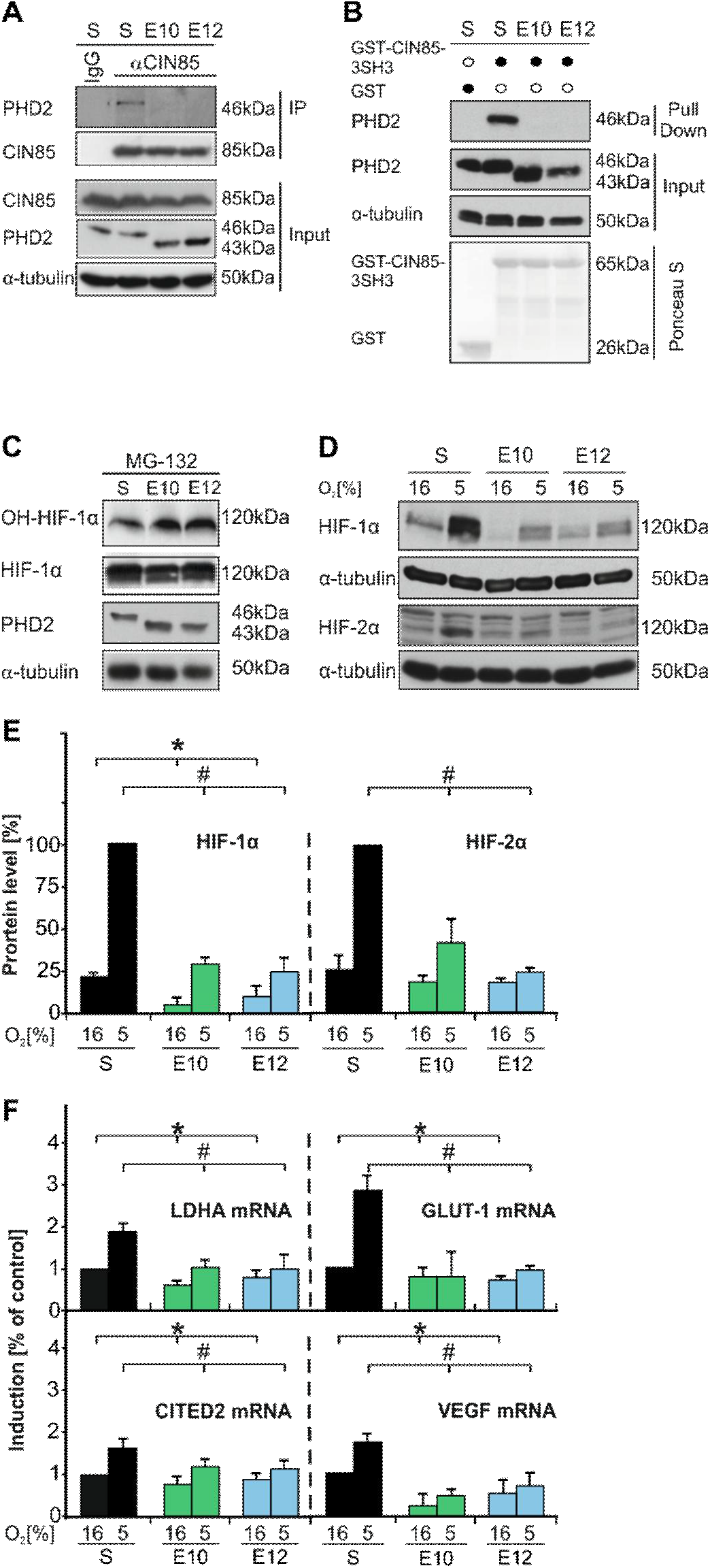
Lack of intracellular CIN85-, PHD2 interaction MDA-MB-231 cells reduces HIF-α levels. (A) IPs were performed with the CIN85 antibody from control S, E10, and E12 cells and probed for PHD2 and CIN85. Blots from the input were probed with CIN85, PHD2, and α-tubulin antibody. (B) WB analysis of PDs using recombinant GST and GST-fusion 3SH3 CIN85 proteins in MDA-MB-231 control S, E10, and E12 cells. Blots from ì PDs were probed with PHD2 antibody; blots from the input were probed with PHD2 and α-tubulin antibodies. Ponceau S stain was used as a loading control. (C) Hydroxy-HIF-1α levels in MG132 treated control S, E10, and E12 cells. Extracts were probed with OH-HIF-1α, PHD2, and α-tubulin antibodies. (D) HIF-1α and HIF-2α levels in control S, E10, and E12 cells cultured under normoxia (16%) or hypoxia (5%) O_2_) for 6 h. Blots were probed with HIF-1α, HIF-2α, and α-tubulin antibodies. (E) Quantification of HIF-1α and HIF-2α levels in control S, E10, and E12 cells, cultured under normoxia or hypoxia. HIF-1α and HIF-2α levels in S cells cultured under hypoxia were set to 100%. Data are mean ±SD (n=4), * significant difference between E10 and E12 cells vs. control (normoxia). # significant difference between E10 and E12 cells vs. control S cells (hypoxia). (F) Quantification of LDHA, GLUT 1, CITED2, and YEGF mRNA levels in control S, E10, and E12 cells cultured under normoxia or hypoxia. The respective mRNA levels in control S cells cultured under normoxia were set to 1. Data are mean ±SD (n=3), * significant difference between E10 and E12 cells vs. control (normoxia), # significant difference between E10 and E12 cells vs. control S cells (hypoxia).

Since our previous data showed that lack of the CIN85-binding domain within PHD2 increases its activity and reduces HIF-1α (Figure 3C-G), we now analyzed whether the lack of the CIN85 binding site in the E10 and E12 cells increased the levels of hydroxy-HIF-1α. The data show that upon inhibition of the proteasomal degradation with MG132 increased levels of hydroxy-HIF-1α could be detected (Figure 4C). Next, we tested whether the edited cells would display reduced protein levels of HIF-1α and HIF-2α under normoxia or hypoxia. Indeed, HIF-1α protein levels were reduced by about 50-60% in E10 and E12 cells compared to control S cells under normoxia and by about 75% under hypoxia (Figure 4D, E). In addition, HIF-2α protein levels were also downregulated in E10 and E12 cells. While there was a trend for decreased HIF-2α levels under normoxic conditions, both E10 and E12 cells showed about 60% lower HIF-2α induction under hypoxia (Figure 4D, E).

In addition, we tested whether the reduction in HIF-α levels has consequences on the expression of HIF-target genes. Indeed, loss of the CIN85-PHD2 interaction in E10 and E12 cells reduced expression of LDHA, GLUT1, VEGF, and CITED2 mRNA under normoxia when compared to S cells and severely reduced their hypoxia-dependent induction (Figure 4F).

It is known, that hypoxia-triggered HIF-α accumulation mainly arises from a PHD-mediated decrease in HIF-degradation, rather than from an increase in mRNA expression (24,27,28). In order to investigate whether the downregulation of HIF-1α and HIF-2α may result from a decrease in mRNA expression, we measured HIF-1α and HIF-2α mRNA expression by qRT-PCR. We did not detect any significant changes in the expression of HIF-1α mRNA between S and E10 or E12 cells (Supplementary Figure 4). However, clone E12 expressed lower levels of HIF-2α mRNA compared to S cells under normoxic conditions (Supplementary Figure 4).

In order to assess whether the absence of the amino acids 75-103 in the engineered cells would lead to a changed expression of the other PHD enzymes we measured the protein levels of PHD1 and PHD3 by Western blot. No differences were found in the expression of PHD1 and PHD3 in E10 or E12 cells (Supplementary Figure 4). In line with the results from the Western blot analyses, E10 and E12 cells did not show changes on the PHD1, PHD2, and PHD3 mRNA levels under normoxic conditions compared to control cells (Supplementary Figure 4). Since the expression levels of PHD1 and PHD3 were not changed in E10 and E12 cells (Supplementary Figure 4), it is possible to conclude that that the observed downregulation in HIF-1α and HIF-2α levels is a result of the altered PHD2. The observed changes cannot be attributed to the off-target effects of the CRISPR/Cas9-mediated *EGLN1* gene editing either (Supplementary Figure 6). Altogether, these data show that the lack of CIN85-PHD2 interaction contributes to a decrease in HIF-α levels.

The above-mentioned findings suggest that the HIF-α levels in the E10 and E12 cells could be resistant to overexpression of CIN85 and that depletion of CIN85 could mimic the loss of the CIN85-PHD2 interaction. To test this, we overexpressed CIN85 in S, E10 and E12 cells. The findings show that overexpression of CIN85 increased HIF-1α and HIF-2α levels only in the S cells but not in E10 and E12 cells (Figure 5A, B). In addition, when we knocked-down CIN85 with shRNA we observed that loss of CIN85 decreased HIFα-levels only in S cells, whereas the already lower HIF-levels in E10 and E12 cells were not affected by the knock-down of CIN85 (Supplementary Figure 5).

**Figure 5.**
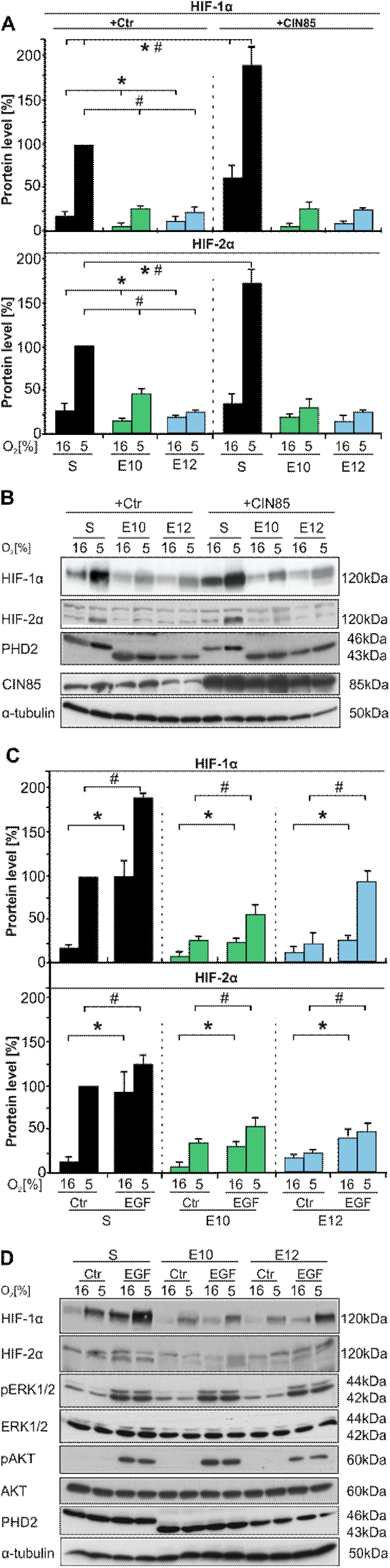
Lack of the CIN85-PHD2 interaction in MDA-MB-231 cells mediates HIF-α resistance to CIN85 overexpression and attenuates EGF-mediated HIF-α induction. (A) Quantification of HIF-1α and HIF-2α levels in MDA-MB-231 control S, E10, and E12 cells transfected with empty vector (+Ctr) and in S, E10, and E12 cells with overexpression of CIN85 (+CIN85). Cells were cultured under normoxia (16%) or hypoxia (5% O_2_) for 4 h. HIF-1α and HIF-2α levels in S (+Ctr) cells under hypoxia were set to 100%. Data are mean ±SD (n=4), * significant difference between E10 and E12 cells vs. S (+Ctr, normoxia); # significant difference between E10 and E12 cells vs. S cells (+Ctr, hypoxia). *# significant differences S (+Ctr) vs. S (+CIN85) under normoxia and hypoxia. (B) WB analysis. Blots were probed with HIF-1α, HIF-2α, PHD2, CIN85, and α-tubulin antibodies. (C) Quantification of HIF-1 α and HIF-2α levels in control S, E10, and E12 cells treated with vehicle (Ctr) or EGF (100 ng/ml). HIF-1α and HIF-2α levels in S (+Ctr) cells under hypoxia were set to 100%. Data are mean ±SD (n=3), * significant difference between vehicle (Ctr) treated S, E10 and E12 cells vs. EGF-treated S, E10 and E12 cells (normoxia); # significant difference between vehicle (Ctr) treated S, E10 and E12 cells vs. EGF-treated S, E10 and E12 cells (hypoxia).

Since CIN85 has a prime function in the activation of ERK1/2 and AKT in the EGF signaling pathway (20), we next investigated whether the loss of the interaction between PHD2 and CIN85 in the E10 and E12 cells affects ERK1/2 and AKT activation and induction of HIF-1α and HIF-2α in response to EGF. The results show that neither basal nor EGF-dependent activation of both ERK1/2 and AKT was altered in E10 and E12 cells when compared to S cells. In addition, EGF mediated an increase in HIF-1α levels in the S cells of about 5-fold under normoxia. EGF was additive under hypoxia and enhanced the hypoxia-dependent induction by about 2-fold. While EGF mediated also an about 5-fold induction of HIF-2α in the S cells under normoxia, no additive effect was seen under hypoxia. Although EGF caused also an induction of both HIF-1α and HIF-2α in E10 and E12 cells under normoxia; this induction was less robust than in the S cells. (Figure 5C, D). Together, these data suggest that the loss of the PHD2-CIN85 interaction is rather selective for HIF-α regulation and does not affect basal and EGF-dependent ERK1/2 and AKT regulation.

### Lack of amino acids 75-103 in PHD2 contributes to less malignant cell properties

MDA-MB-231 is a triple negative breast cancer cell line known to have an aggressive metastatic phenotype. Therefore, we further investigated whether the expression of Δ75-103 PHD2, and thus the lack of CIN85-PHD2 interaction, contributed to any changes in morphology and malignancy of these cells. In order to visualize the cell shape and surface composition, we imaged S, E10, and E12 cells by scanning electron microscopy (SEM) (Supplementary Figure 6A). Cells expressing Δ75-103 PHD2 were elongated with fewer protrusions on the leading edge of the cells. Additionally, we investigated whether the absence of amino acids 75-103 had an impact on the intracellular localization of PHD2. Immunofluorescence revealed that Δ75-103 PHD2 did not change the cellular localization. The distribution was mainly cytoplasmic, which was similar to the wt PHD2 in S control cells. Visualization of α-tubulin in S, E10, and E12 further verified the altered shape and morphology of E10 and E12 cells, in line with the results of the electron microscopy (Supplementary Figure 6B). After observing the changes in cellular shape in E10 and E12 cells, we assessed cellular proliferation, the ability to form colonies, the motility of S, E10, and E12 cells, and tumor forming potential by injecting these cells into nude mice.

When performing live-cell proliferation analyses using the IncuCyte^®^ ZOOM System we found that proliferation of the two clones E10, and E12 expressing Δ75-103 PHD2 was slower than that of MDA-MB-231 S control cells (Figure 6A). While it took 72 h for the control S cells to reach 100% confluence, E10 cells were able to display the same confluence level only after 108 h, whereas E12 cells showed only 80% confluence at the end of a 117 h experiment (Figure 6A).

**Figure 6.**
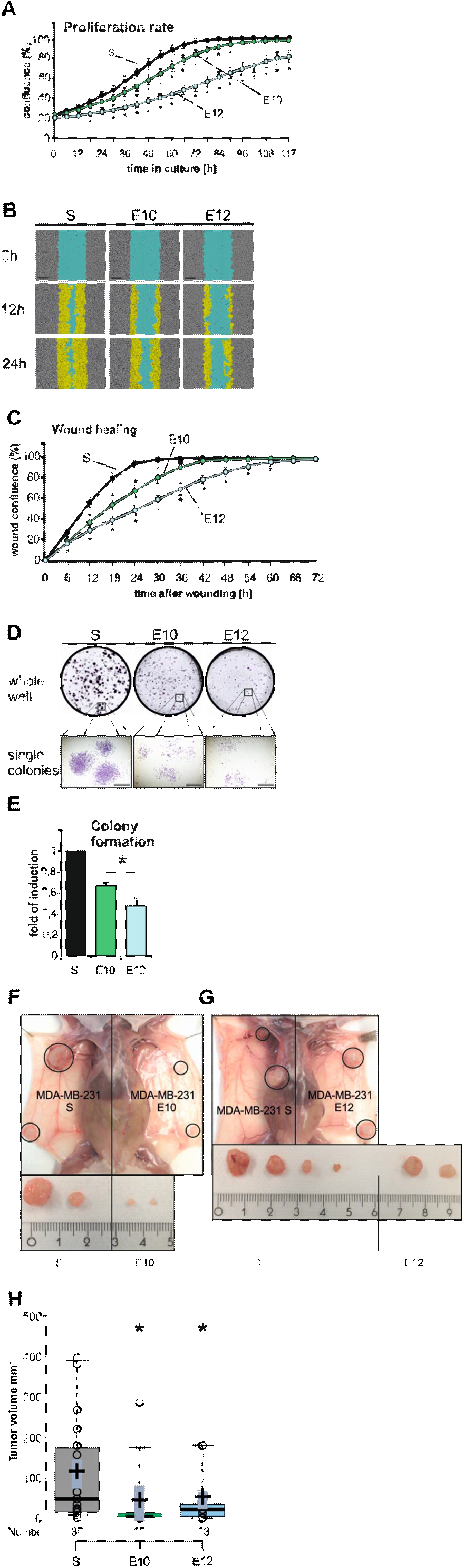
Lack of amino acids 75-103 in PHD2 reduces malignant properties of MDA-MB-231 cells. (A) Live cell proliferation analysis of MDA-MB-231 control S, E10, and E12 cells (cf Material and Methods). Data are mean ±SD (n=5), * significant difference between relative confluence values of Δ75-103 PHD2 expressing cells (E10 and E12) vs. control S cells at each time point. (B) Representative images of the wound closure of control S, E10, and E12 cells at 0, 12 h and 24 h time point after introduction of the wound. Scale bar 300 um. (C) Live cell wound closure analysis of MDA-MB-231 S, E10, and E12 cells (cf Material and Methods). Data are mean ±SD (n=4), * significant difference between wound confluence values of Δ75-103 PHD2 expressing cells (E10 and E12) vs. control S cells at each time point. (D) Representative images of the whole cell culture wells and single colonies formed by MDA-MB-231 S, E10, and E12 cells. Scale bar 1 mm. (E) Quantification of the colony formation assay. The total number of colonies per well was set to 1 in control S cells. Data are mean ±SD (n=4), * significant difference between the number of colonies formed by E10 and E12 cells vs. control S cells. (F,G) MDA-MB-231 control S, E10, and E12 cells were injected into the thoracic and inguinal mammary fat pads of female athymic nude mice (6 animals per group). Representative image of tumors derived from control S vs. E10 (F) and control S vs. E12 (G). (H) Box plot indicating the respective volume differences of the control S-derived tumors compared to the E10- and E12-derived tumors. The center lines show the medians; crosses represent sample means; grey bars within boxes indicate 83% confidence intervals of the means; the box limits indicate the 25th and 75th percentiles as determined by R software; whiskers extend to minimum and maximum values; data points are plotted as open circles. Number = 30, 10, 13 tumor sample points. * significant difference of the means S- vs. E10- and E12.

Additionally, we evaluated cellular motility in a low-serum wound (scratch) assay. Confluent MDA-MB-231 control S, E10, and E12 cells were wounded and wound closure was monitored for 72 h. In line with the data from proliferation assays, both of the clones expressing Δ75-103 PHD2 were almost 50% less motile than the control cells during wound closure. While the control S cells were able to close the scratch area in about 24-30 h, it took E10 and E12 cells about 12-18 h and 26-32 h longer, respectively (Figure 6B, C).

The ability to grow and form colonies from single cells is a very important malignant property of cancer cells. Therefore, we performed colony formation assays by plating cells at very low density and allowing them to grow for 10 days. Colony formation was drastically impaired in E10 and E12 cells (Figure 6C, D). The number and size of colonies formed by control S cells was higher than in E10 and E12 cells. In addition, the colonies of control cells were dense and displayed an overlapping growth pattern, while E10 and E12 cells were disseminated within the colony (Figure 6C, D).

We next tested whether the observed reduced proliferation and impaired migration of E10 and E12 cells are the result of the increased PHD2 activity and reduced HIF-1α levels. To do this, we treated the cells with the HIF prolylhydroxylase inhibitor FG-4592 (Roxadustat), or transfected the cells with either HIF-1α or PHD2 expression vectors. Treatment of cells with FG-4592 stabilized HIF-1α and counteracted the reduced proliferation and migration in E10 and E12 cells without having significant effects in control S cells (Supplementary Figure 7A-G). Although overexpression of HIF-1α promoted proliferation as well as migration in all cells, its action was more pronounced in E10 and E12 cells. Like with FG-4592, this abolished the differences in proliferation and wound healing in E10 cells and largely also in E12 cells (Supplementary Figure 7A-G). By contrast, overexpression of full length PHD2 inhibited proliferation and wound healing significantly only in S cells and marginally but insignificantly in E10 and E12 cells (Supplementary Figure 7A-G). Together, these findings support the view that the observed cellular phenotypes of E10 and E12 cells are largely dependent on both, the activity of PHD2 and HIF-1α.

Since the above-mentioned assays pointed to a less malignant phenotype of E10 and E12 cells compared to control S cells, we next investigated whether these characteristics are also present *in vivo*. Therefore, we transplanted MDA-MB-231 control S vs. E10 and S vs. E12 cells, into the thoracic and inguinal mammary fat pads of 18 female athymic immune deficient nude mice and followed tumor growth by palpation and size measurement for up to eight weeks. Although all cells were able to form tumors in these mice, it was obvious that the arisen tumors from control S cells were of bigger volume, compared to tumors formed from E10 and E12 cells (Figure 6F-G). While the average volume of tumors from control S cells was about 138 mm^3^ at the time of necropsy, the E10-derived and E12-tumors had a volume of about 36 mm^3^ and 44 mm^3^, respectively (Figure 6F-H). We also noted that the E10 and E12 cell-derived tumors were paler when compared to the tumors formed from control S cells, which is likely a result from reduced angiogenesis due to lower HIF-α levels (Figure 6F-H).

Collectively, these data show that MDA-MB-231 cells expressing Δ75-103 PHD2, and thus lacking the interaction between PHD2 and CIN85, are less motile, have a lower proliferative, as well as tumor forming potential.

## Discussion

The current investigation shows that the pro-oncogenic adaptor protein CIN85 is a novel inhibitory binding partner of PHD2 (Figure 1). We defined the nature of CIN85-PHD2 complex formation by revealing the importance of the three N-terminal SH3 domains of CIN85 and the proline and arginine residues within the N-terminus of PHD2, lying outside of the Zn-finger domain of PHD2 (Figure 2-3). By performing CRISPR/Cas9-mediated *EGLN1* gene editing, we created MDA-MB-231 cell lines, expressing a PHD2 lacking amino acids 75-103, which was unable to bind CIN85 (Figure 4). The abrogation of the PHD2-CIN85 interaction led to lower HIF-1α and HIF-2α levels (Figure 4-5), thus contributing to a less malignant phenotype of MDA-MB-231 cells (Figure 6).

The overexpression of the EGFR-adaptor protein CIN85 is known to influence the development and progression of various types of cancers, among which are carcinomas of the head and neck (21), cervix (29), and colon (22). Our own research showed that CIN85 is highly overexpressed in breast carcinoma (20), and contributes to breast cancer pathogenesis via the attenuation of EGFR and ErbB2/HER2-downregulation (14,20,30). In addition, the mechanisms behind CIN85-driven carcinogenesis include an enhancement of transforming growth factor-β (TGF-β) signaling (31), promotion of cell invasiveness via associations with Mucin 1 (MUC1) (32), diverse cytoskeletal elements, and focal adhesion kinase (FAK) activation (33,34). Moreover, our previous findings indicated that CIN85 contributes to cancer progression via stabilization of HIF-1/2α (as well as the expression of HIF-target genes such as PAI-1 (23). The current study extended these findings and verified that the direct interaction of CIN85 and PHD2 is responsible for inhibition of the PHD2-mediated prolyl-hydroxylation of HIF-α subunits.

The HIF family of transcription factors are essential molecular regulators of normal development (35) and at the same time serve as crucial contributors to the progression of various types of cancers (36,37). Although HIF-α levels are regulated via multiple mechanisms (6,7), the hydroxylation-dependent mechanism is considered to have a major impact on HIF-1α-degradation. All members of the HIF-PHD family are able to hydroxylate HIF-1α *in vitro* (38–40); however, PHD2 is found to be the most important hydroxylase to regulate HIF-1α levels under normoxic conditions (24,28). The activity of PHD2 is critical for HIF-α stability; however, alterations in the enzymatic function of PHD2 are not always caused by the lack of oxygen (41). The enzymatic function of PHD2 can be indirectly altered through the association with other proteins that interfere with the PHD2/HIF-αs/VHL complex formation.

The majority of PHD2 interactors contribute to the downregulation of HIF-α levels. For example, inhibitor of growth family member 4 (ING4) (42), p23 from the HSP90-machinery (43), and the runt-domain transcription factor (44) were reported to promote the association between PHD2 and HIF-α that leads to a faster downregulation of HIF-1α (45). Another example is the direct interaction of the adaptor proteins LIMD1 and RHO-related BTB domain-containing protein 3 (RHOBTB3) with PHD2 and VHL simultaneously (46). The association of PHD2 with these adaptors promotes the assembly of hydroxylation-dependent degradation machinery, which further contributes to lower levels of both HIF-1α and HIF-2α (46,47). Since decreased levels of HIF-αs are usually considered to favor patient survival, the participation of RUNX3 (48), ING4 (49), LIMD1 (50), and RHOBTB3 (47) in downregulating HIF-αs additionally confirms the known tumor-suppressing role of these proteins in human cancers. Interestingly, our data show that the strength of the interaction between PHD2 and CIN85 varies between the cell lines investigated. This could suggest that (an)other factor(s), existence of which could vary between the different cell types, may contribute to the strength of the interaction and future studies will show whether this is the case.

The results of our study indicate that the interaction of CIN85 with PHD2 leads to a very different outcome by contributing to the increased stability of HIF-1α and HIF-2α. Our current investigation describes a direct interaction with PHD2 as a mechanism by which CIN85 regulates HIF-1α and HIF-2α stability in breast cancer cells. This event appears to occur in the cytoplasm since we found that the CIN85-PHD2 interaction was restricted to the cytoplasm (Figure 1). This is in line with reports that both PHD2 and CIN85 are cytoplasmic proteins (Take, Watanabe et al. 2000), although PHD2 has also been found in the nucleus (51). In addition, we showed that CIN85 utilizes three SH3 domains to facilitate the binding to PHD2 (Figure 2).

SH3 domains are non-catalytic domains able to bind proline-rich sequences in a large number of proteins (52,53). While the majority of SH3 domains bind ‘typical’ proline-rich motifs like PXXP (where X is any amino acid), the SH3 domains of CIN85 are reported to recognize atypical proline-arginine sequences (54). By using a variety of PHD2 and CIN85 constructs in immunoprecipitation and GST-pull down experiments, we found that a proline-arginine (PR)-rich area between amino acids 77-100 is the most important for the direct interaction with CIN85 (Figure 1-3). While PHD2-binding partners are known to utilize either the Zn-MYND finger domain (55,56), or the catalytic domain of PHD2, we define CIN85 as the first interactor recognizing a different binding motif in the sequence of PHD2 (Figure 3). The mapping of the binding motif for CIN85 not only identified critical residues mediating the CIN85-PHD2 complex formation, but also revealed the existence of a functional sequence in the N-terminus of PHD2 able to mediate protein-protein interactions. Although this approach was straight forward, there exists also the possibility that the systematic mutation of the proline residues may have led to a conformational change in PHD2 in other regions, thus influencing the binding between CIN85 and PHD2 indirectly. Interestingly, the PHD2 variants lacking the ability to interact with CIN85 showed higher catalytic activity in the *in vitro* hydroxylation assay, thus leading to lower levels of HIF-1α. Although the structural basis for oxygen degradation domain selectivity of the catalytic PHD regions are quite well known(57)), the crystal structural details of the PHD2 N-terminus harboring the CIN85 interaction site remain so far unresolved. However, the current data are in line with earlier findings showing that the N-terminus of PHD2 is inhibitory (58). This, in line with the in vitro hydroxylation assays of this study, suggests that binding of CIN85 might regulate PHD2 activity at the N-terminus by reducing the accessibility of its catalytic domain to a substrate.

After establishing the binding pattern between CIN85 and PHD2, we created a cell system lacking this molecular interplay using CRISPR/Cas9-mediated *EGLN1* gene editing. Two independent cell clones expressing PHD2 lacking amino acids 75-103 and consequently the interaction between CIN85 and PHD2 were generated (Figure 4). The MDA-MB-231 Δ75-103 PHD2-expressing cell lines E10 and E12 showed, as expected, lower HIF-1α and HIF-2α protein levels (Figure 5), and slower proliferation, decreased motility, as well as a reduced tumor forming potential in nude mice when compared to the respective control cells (Figure 6).

Our previous findings indicated that overexpression of CIN85 stabilizes HIF-1α levels, and notably, that downregulation of CIN85 reversed this effect (23). The results of the current study allow us to conclude that lower HIF-1α levels under normoxic conditions in Δ75-103 PHD2 expressing cells originate from decreased protein stability and cannot be attributed to the differences in HIF-α mRNA expression. In addition, we can associate the observed downregulation in HIF-1α and HIF-2α levels with Δ75-103 PHD2 function, since the expression of the two other PHDs (PHD1 and PHD3) was not affected (Supplementary Figure 4).

Altogether, our study is the first to describe the relation between PHD2 and CIN85. We identified CIN85 as a novel inhibitory binding partner of PHD2, able to increase HIF-α levels and, thus, to promote HIF-α stability and breast cancer malignancy.

## Materials and methods

### Chemicals

All biochemicals and enzymes were of analytical grade and were purchased from commercial suppliers. EGF was from Sigma-Aldrich, restriction enzymes were from Thermo Scientific, FG-4592 was from Cayman Chemical.

### Cell culture

Human embryonic kidney 293 (HEK-293, # CRL-1573), human breast carcinoma cell lines (MDA-MB-231 (#HTB-26), Hs578T (#HTB-126) and BT-549 (#HTB-122) were purchased from ATCC. HEK-293, MDA-MB-231, and Hs578TH were maintained in Dulbecco’s modified Eagle medium (DMEM) supplemented with 10% fetal calf serum (FCS), 2 mM L-glutamine, 50 IU/ml penicillin, and 50 μg/ml streptomycin in a humidified atmosphere containing 5% CO_2_, 16% O_2_, 79% N_2_ at 37°C. The cell line BT-549 was maintained in Roswell Park Memorial Institute-developed (RPMI) 1640 medium. When indicated, the cells were incubated under hypoxic conditions in a Ruskinn Sci-Tive-N hypoxia workstation under 5% or 1% O_2_, 5% CO_2_ balanced with N_2_ for 6 h. All cell lines underwent mycoplasma testing before use.

### Plasmids and site directed mutagenesis

pClneo-Myc-CIN85 D111G was a gift from Yutaka Hata (Addgene plasmid # 47935) (59), the D114G mutation was eliminated by site directed mutagenesis (QuickChange mutagenesis kit, Promega). The constructs for the bimolecular fluorescence complementation assay (BiFC) were generated by PCR, the respective PCR products were subcloned into the BamHI and XbaI sites (for PHD2) or EcoRI and XbaI sites (for CIN85) of pcDNA3-YN (non-fluorescent N-terminus of YFP) or pcDNA3-YC (non-fluorescent C-terminus of YFP) plasmids, respectively (60). The constructs encoding CIN85 deletion variants and the ones allowing the expression of the recombinant GST-SH3A, GST-SH3B, GST-SH3C, GST-Pro, GST-Ser, and GST-CC fusion proteins were previously described (61,62).

The expression vectors for PHD2, PHD1 and PHD3 were described (63) PHD2 deletion mutants lacking amino acids 1-26 (Δ1-26 PHD2), 10-58 (Δ10-58 PHD2), 52-98 (Δ52-98 PHD2) and 99-136 (Δ99-136 PHD2) were generated by site directed mutagenesis of pcDNA3.1-PHD2-V5-6xHis creating additional BamHI (Δ1-26 PHD2), SgsI (Δ52-98 PHD2, Δ99-136 PHD2), and SfiI (Δ10-58 PHD2) restriction sites in the coding sequence of *EGLN1* (PHD2). Afterwards, mutants were digested with BamHI, SgsI, SfiI, and XbaI (Thermo Fisher Scientific) respectively, fragments were gel-purified (Gel/PCR DNA fragments extraction kit, GeneAid), re-ligated, and transformed into *E.coli* XL-1 blue competent cells. PHD2 deletion mutants lacking amino acids 1-136 (Δ1-136 PHD2) and 210-426 (Δ209-426 PHD2) were described previously (Kozlova, Wottawa et al. 2016).

The plasmids expressing the point-mutated PHD2 variants, P77P79P80P84P85P86A-PHD2 and P86P87R91P93R94A-PHD2, were obtained via subcloning of synthetically produced DNA fragments (Eurofins Genomics) between BamHI and SgsI restriction sites of pcDNA3.1-PHD2-V5-6xHis. The construct P86P87R91P93P94R99R100A-PHD2 was obtained by site-directed mutagenesis of pcDNA3.1-P86P87R91P93R94A-PHD2-V5-His.

The constructs allowing the expression of recombinant GST-fusion PHD2 proteins were generated by PCR, and the respective PCR products were subcloned into the BamHI and EcoRI sites of pGEX-5X-1 (GE Healthcare). The psPAX2 was a gift from Didier Trono (Addgene plasmid # 12260) and the pVSVg was a gift from Robert Weinberg (Addgene plasmid # 8454) (64). LentiCas9-Blast and lentiGuide-Puro were a gift from Feng Zhang (Addgene plasmids #52962 and #52963, respectively) (65,66). The lentiGuide-Puro vectors were further modified by subcloning of the corresponding annealed oligonucleotides with BsmBI overhangs. All constructs were verified by DNA sequencing. Primers used in the study are listed in Supplementary Table 1.

### Protein preparation and Western blotting

The cells were lysed in lysis buffer [50 mM Tris–HCl, pH 7.5, 150 mM NaCl, 1% Triton X-100, 1 mM o-vanadate, 50 mM NaF, 2 mM EDTA, 1 mM PMSF, complete protease inhibitor cocktail tablet (Roche)], kept on ice for 10 min and centrifuged at 14 000 g for 20 min at 4°C (67). For EGF stimulation cells were cultured in starvation medium (DMEM containing 0.1% FBS) 24 h, then treated with EGF (100 ng/ml) and lysed as described above. The levels of HIF-1α, HIF-2α, PHD1, PHD2, PHD3, Myc-CIN85, V5-PHD1, V5-PHD2, V5-PHD3,Flag-CIN85, CIN85, phospho-AKT, and phospho-ERK1/2 were detected by Western blotting from whole cell extracts. Proteins (20-100 μg per sample) were separated by electrophoresis on 7.5-12% polyacrylamide gels and transferred to nitrocellulose membranes. The membranes were incubated with the following antibodies: HIF-1α (#610959, BD Biosciences), HIF-2α (#NB100-122, Novus Biologicals), PHD1 (NB100-310, Novus Biologicals), PHD2 (#3293, Cell Signaling), PHD3 (NB100-139, Novus Biologicals), Myc-Tag (#2278, Cell Signaling), V5-Tag (#R96025, Thermo Fisher), Flag M2 (#F1804, Sigma-Aldrich,), CIN85 (?), phospho-AKT (pSer473) (#9271, Cell Signaling), AKT (#9272, Cell Signaling), phospho-ERK1/2 (pThr202/ pTyr204) (#9101, Cell Signaling), ERK1/2 (#9107, Cell Signaling), α-tubulin (B-5-1-2) (#T5168, Sigma-Aldrich), and β-actin (A5316, Sigma-Aldrich) overnight at 4°C. Appropriate secondary antibodies (peroxidase-conjugated IgG (Biorad)) were used. The ECL kit (GE Healthcare) was used for signal detection. Blots were quantified by densitometry with the Image Quant TL program (GE Healthcare); densitometry data were normalized to α-tubulin or β-actin.

### Immunoprecipitation

For the co-immunoprecipitation of CIN85 and PHDs, HEK-293 cells were transiently transfected with expression plasmids encoding Myc-tagged CIN85 and V5-tagged PHDs (PHD1, PHD2, and PHD3) or Flag-tagged CIN85 lacking either SH3A, SH3B, SH3C or all three SH3 domains, and V5-tagged PHD2. Immunoprecipitations were carried out as described (6). Cells were harvested 48 h post-transfection, washed twice with ice-cold phosphate buffered saline (PBS) and lysed as described above. Aliquots of cleared HEK-293 cell lysates containing 1 mg of total protein were mixed with protein G Sepharose beads (GE Healthcare) and Myc-tagged CIN85 was immunoprecipitated with the Myc-tag antibody; the Flag-tagged CIN85 variants were immunoprecipitated with the Flag M2 antibody at 4°C overnight. For the co-immunoprecipitation of CIN85 and PHD2 at the endogenous level, CIN85 was immunoprecipitated from the lysates of MDA-MB-231, Hs 578T, and BT-549 cells with the SH3A-CIN85 monoclonal antibody (68). The next day the beads were washed 5 times with lysis buffer, the immune complexes were then resolved by SDS-PAGE and analyzed with antibodies against the Myc and V5 epitope, or PHD2.

### GST-pull down

Recombinant GST-fusion proteins containing various fragments of CIN85 were expressed in *E.coli* BL-21, while the GST-fusion PHD2 proteins were expressed in Rosetta 2 DE3 pLys. The recombinant proteins were captured from bacterial lysates via glutathione-sepharose beads (GE Healthcare) according to the manufacturer’s protocol. For the pull down assays HEK-293 cells were transiently transfected with expression plasmids encoding V5-tagged PHD2 or Myc-tagged CIN85. The aliquots of lysates were mixed with either GST or GST-fusion proteins immobilized on glutathione sepharose beads overnight at 4°C. For investigation of the binding of endogenous PHD2 to GST-3SH3-CIN85, cell lysates from MDA-MB-231 cells were used. Bound immune complexes were collected and analyzed as above with antibodies against the Myc and V5 epitope, or PHD2. The nitrocellulose membranes were stained with Ponceau S to visualize the equal loading of GST-fusion proteins used for the reaction.

### Fluorescence microscopy

For visualization of the BiFC signal, BT-549 cells were plated on glass coverslips and transfected with the following combination of constructs: 1). pcDNA3-CIN85-YN + pcDNA3-PHD2-YC, and 2). pcDNA3-YN + pcDNA3-YC (negative control), while transfection with pcDNA3-YFP alone served as a positive control. 48 h post-transfection the cells were fixed with 4% paraformaldehyde (PFA). The coverslips were washed and stained with bisbenzimidine (Hoechst stain, Sigma-Aldrich) to visualize the nuclei of the cells. Next, coverslips were washed in PBS and in water, and mounted using Shandon Immumount mounting media (#9990402, Thermo Fisher). Confocal microscopy was performed using a Zeiss Observer Z1 equipped with a LSM 700 confocal unit, 63x PlanApo oil immersion objective and appropriate filter sets for Hoechst 405, and YFP, and Zen2009 software. To visualize the localization of PHD2 in MDA-MB-231 cells after CRISPR/Cas9-mediated *EGLN1* (PHD2) gene editing, the cells were plated on glass coverslips. The next day the cells were rinsed with PBS, fixed with 4% PFA, kept in blocking buffer (1xPBS, 1% BSA, 0.1% Saponin) for 60 min and further incubated with rabbit PHD2 (NB100-138 Novus Biologicals) and mouse α-tubulin primary antibodies for 60 min. The coverslips were washed and incubated with the corresponding Alexa Fluor 546 and Alexa Fluor 488 conjugated secondary antibodies in a 1:500 dilution (#A-11035, #A-11001, Thermo Fisher Scientific) at RT for 60 min. Then, the coverslips were examined by confocal microscopy using appropriate filter sets for Hoechst 405, Alexa Fluor 488, Alexa Fluor 546, and Zen2009 software.

### CRISPR/Cas9-mediated EGLN1 gene editing

To obtain a cell line expressing PHD2 lacking the critical amino acids for the CIN85-PHD2 interaction, the clustered regularly-interspaced short palindromic repeats (CRISPR)/Cas9 approach was used (Ran, Hsu et al. 2013). The sequence corresponding to the nucleotides 3329-3576 of the first exon of *EGLN1* was submitted as a template to the mit.crispr.edu online tool. Two target sequences, allowing the introduction of two proximal double-strand breaks (DSBs) in the EGLN1 gene were chosen and referred to as sgRNA-EGLN1-219 and sgRNA-EGLN1-292. Two missense sequences (targeting bacterial *lacZ* and a scrambled sequence OG from OriGene) were chosen as corresponding negative controls. In order to disable the repair of DSBs by non-homologous end-joining (NHEJ), after the introduction of Cas9-sgRNAs, the cells were treated with the DNA ligase IV inhibitor Scr7 (Cayman Chemical) (69,70). The sequences of the oligonucleotides used are listed in Supplementary Table 1.

### Lentivirus-mediated expression of sgRNAs and Cas9

For the generation of lentiviruses, 70-80% confluent HEK-293T cells were co-transfected with psPAX2, pVSVg, and one of the following vectors: lentiCas9-Blast, lentiGuide-Puro EGLN1-sgRNA-219, lentiGuide-Puro EGLN1-sgRNA-292, lentiGuide-Puro SCR1-sgRNA-OG, or lentiGuide-Puro SCR2-sgRNA-LacZ by Lipofectamine 2000 (Thermo Fisher Scientific). Media was collected over a period of 24 h to 72 h post-transfection in 24 h batches, pooled, centrifuged and filtered through a 0.45 μm filter. As a first step, MDA-MB-231 cells were infected with the virus, which allowed the expression of Cas9 endonuclease. The next day virus-containing media was replaced with fresh media, which allowed the cells to recover. 48 h post-infection, the cells were trypsinized and re-seeded in the presence of Blasticidin (30 mg/ml) followed by 7 days of selection. Next, Cas9-expressing MDA-MB-231 cells were infected with the combination of the two sgRNAs, either targeting the *EGLN1* locus (EGLN1-sgRNA-219 and EGLN1-sgRNA-292), or scrambled sgRNAs (SCR1-sgRNA-OG and SCR2-sgRNA-LacZ). Similarly, the cells were allowed to recover, then they were cultured in the presence of Puromycin (1 mg/ml) for 24 h and the DNA ligase IV inhibitor Scr7 (1 μM) for 72 h. Afterwards, clonal cell populations were generated by fluorescence activated cell sorting (FACS).

### Genomic DNA extraction and genotyping

Total DNA was extracted from the cells using the Quickextract DNA Extraction solution (Epicentre) (#QE0905T, Immuno Diagnostics) according to the manufacturer’s instruction. Later the DNA was used for PCR with Herculase II fusion polymerase (#600677 Agilent Technologies) and a set of genotyping primers (Egln1-2739F and Egln1-3576R) listed in Supplementary Table 1, which allowed the amplification of a 908 bp region from wt *EGLN1*, and 821 bp in EGLN1-edited cells. In order to better observe the size-shift and potential presence of homozygous deletions, an 8.5% native PAGE-based genotyping approach was used (71) that allowed visualization of the presence of either edited or wt *EGLN1* alleles due to the formation of DNA heteroduplexes. The obtained PCR-products were separated by PAGE and visualized by ethidium bromide staining. Screening for the potential off-target effects was performed via PCR amplification of the potential DSBs in the exonic area of the three genes for EGLN1-sgRNA-219 and EGLN1-sgRNA-292. In all cases, the respective PCR products were electrophoresed on agarose gels, gel-purified, and sequenced using the same forward and/or reverse primer.

### RNA extraction and qRT-PCR

Total RNA was isolated from cells with a GenElute mammalian total RNA miniprep kit (Sigma-Aldrich). Reverse transcription (RT) was performed with 1 μg RNA using the first-strand cDNA synthesis kit (Quanta Bioscience, GE Healthcare). As a verification of the CRISPR/Cas9-mediated *EGLN1* editing, cDNA was used in a PCR reaction using the same set of genotyping primers as for the gDNA genotyping; subsequently the purified PCR products were sequenced using the same forward and/or reverse primer. qRT-PCR was performed with cDNA diluted 1:25 and the iTaq Universal SYBR Green Supermix reaction kit (Biorad) in combination with the Applied Biosystems 7500 thermal cycler (Applied Biosystems). HIF-1α, HIF-2α, PHD1, PHD2, PHD3, GLUT1, LDHA, CITED2, and VEGF-A relative mRNA expression was determined using the ΔΔCt data analysis method (72). Ribosomal 18S RNA and beta-actin (*ACTB*) were used as housekeeping genes (73). The primers used are listed in Supplementary Table 1.

### Scanning electron microscopy

Cells were grown on round glass coverslips at low density, washed with PBS and fixed with 2.5% glutaraldehyde in 0.1 M phosphate buffer for 30 min, washed with 0.1 M phosphate buffer and imaged in the Tissue Imaging Core Facility (Biocenter Oulu) with a Sigma HD VP FE-SEM equipped with ET-SE and In-lens SE detectors as described (74).

### Monolayer colony formation assay

Cells were seeded onto 6 well plates (5×10^2^ cells/well) and grown for 10 days. Afterwards, the cells were rinsed with PBS, fixed with 4% PFA, stained with 1% crystal violet solution for 30 min and extensively washed. After air-drying, the plates were photographed and colonies counted using ImageJ software. Single colonies were later imaged with a Leica MZFLIII microscope.

### Live Cell Imaging Assays

For real-time quantitative live-cell proliferation analysis, cells were seeded onto 96-well plates (5×10^3^ cells per well) in full media. The next day the plate was moved into the IncuCyte^®^ ZOOM System (Essen BioScience) for live phase contrast recording of cell confluence for 117 h in 3 h intervals. For the scratch wound assay, cells were seeded onto 96-well Essen ImageLock Plates (Essen Bioscience) (4×10^4^ cells per well with 1% serum). The following day the confluent cell monolayer was wounded with the 96 PTFE pin Wound Maker (Essen Bioscience). The live wound closure was phase contrast-imaged for 72 h in 3 h intervals. In both assays, the confluence analysis was performed using the basic IncuCyte software settings.

### In vitro hydroxylation assay

The catalysts (PHD2 wt, Δ52-98 PHD2, and PR mut 2) in pcDNA vectors containing a T7 promoter were were *in vitro* transcribed and translated (IVTTed) in rabbit reticulocyte lysate by using the TnT^®^ Quick Coupled Transcription/Translation System (Promega) in the presence of [^35^S]Met. An aliquot of the IVTTed catalysts was analyzed on a 10% SDS-Page gel to ensure equal expression levels. The rest of the IVTTed catalysts (45 ul) was used to in vitro hydroxylate [^3^H]Pro-labelled HIF-1α-ODDD in the presence of cofactors Fe^2^+ (5 uM), 2-oxoglutarate (320 uM) and ascorbate (2 mM) at 37°C for 30 min. The generation of [^3^H] 4-hydroxyproline was determined by a radiochemical assay (for details see (38,75).

### Surface plasmon resonance (SPR)

SPR binding studies were performed using a Biacore T200 biosensor system (GE Healthcare, Uppsala, Sweden). All the SPR-based materials were acquired from GE Healthcare. Human PHD2 and CIN85 were produced in insect cells and purified as described earlier (38,75). PHD2 was coupled to a CM5 chip using an amine coupling kit according to the instructions of the manufacturer at pH 5.5, at a level of 4000 RU. Sensor preparation and interaction analyses were performed at 25 °C in a PBS-P [10 mM phosphate buffer, 150 mM NaCl, 2.7 mM KCl, 0.05% P20 (pH 7.4)] at a flow rate of 30 μl/min for 2 min, and a 10-min dissociation phase followed each injection. Before the next injection cycle, the surface was regenerated with a 60-s injection of 1 M NaCl and allowed to stabilize for another 60 s. The BIAevaluation software version 2.0 (Biacore AB) was used to analyse the data. Bulk effects were subtracted using a reference surface without PHD2. Furthermore, a sensorgram of the injection of buffer alone was subtracted from the data.

### Xenograft mouse model

5×10^5^ cells of CRISPR/Cas9-generated MDA-MB-231 cells expressing edited *EGLN1* (PHD2) (E10 and E12) together with scrambled control cells (S) were injected in the thoracic and inguinal mammary fat pad of 4 weeks old female athymic nude mice (Envigo, former Harlan). The mice were housed in IVC cages with water and food *ad libitum* for up to five weeks. The animals were euthanized by CO_2_ inhalation and additional cervical dislocation, and the tumors were collected. All animals were housed in the laboratory Animal Center of the University of Oulu in specific pathogen-free facilities on a 12 h light/dark cycle, at a constant temperature of 22 °C. The protocol for animal use and experiments was approved by the National Animal Experimental Board of Finland (ELLA) as well as the Animal Welfare Body of the Laboratory Animal Center and conducted according to the EU directive 2010/63/EU. The volume of the tumors formed was calculated using the ellipsoid volume formula as described (76).

### Statistical analysis

The results are presented as mean values ± SD of at least 3 independent experiments. Statistical analyses were performed using Student’s two-tailed t-test. Differences of p ≤ 0.05 were considered statistically significant.

## Supplementary material summary

Supplementary material consists of 1 table listing the sequences of the oligonucleotides used in the study followed by 6 supplementary figures containing the results of the GST-pull downs between CIN85 and PHD2 fragments (Supplementary Figure 1 and 2), generation of E10 and E12 MDA-MB-231 cells lacking the CIN85-PHD2 interaction part of PHD2 (Supplementary Figure 3), Proliferation and wound healing assay data from CRISPR/Cas9 edited *EGLN1* cells in the presence of FG-4592, overexpressed HIF-1α and PHD2 (Supplementary Figure 7), Western Blot data and RT-qPCR from CRISPR/Cas9 edited *EGLN1* cells (Supplementary Figure 5 and 6), and analyses of the off-target effects in the CRISPR/Cas9 edited *EGLN1* cells (Supplementary Figure 7).

## Acknowledgments

The authors are grateful to Dr. Svetlana Marchenko for help with the GST-pull down experiments in the beginning of the study, to Dr. Kristian Koski for help with protein purification, Jaana Träskelin and Dr. Satu Myllymäki for help with the production of lentiviral particles, and to Lea Boten, Jonas Böhm, Lea Cleve, Anabel Arnemann (School of Life Science Hamburg, Germany) and Maire Jarva for excellent technical assistance.

## Author contribution

N.K. planned the study, designed the experimental work, performed the experimental work, analyzed the data, and wrote the manuscript; D.M. performed experimental work and assisted with the animal work; A.S. planned, performed experimental work and analyzed the data in the beginning of the study; P.K. performed the *in vitro* hydroxylation assay and analyzed the data; E.B. assisted with the experimental work with recombinant proteins; E.Y.D. assisted with the design of CRISPR/Cas9-mediated *EGLN1* editing work, K.R. assisted with the animal and cell culture work; A.H and S.K. assisted with BiFC experiments; A.M. assisted with the design of lentiviral work; I.M. performed the SEM; V.G. performed cell sorting; L.R. assisted with the experimental design of the protein work; L.D. planned, designed the experimental work and analyzed the data in the beginning of the study; T.K. planned the study, supervised the study, took part in data analysis and writing of the manuscript.

## References

(1) Laughner E, Taghavi P, Chiles K, Mahon PC, Semenza GL. HER2 (neu) signaling increases the rate of hypoxia-inducible factor 1alpha (HIF-1alpha) synthesis: novel mechanism for HIF-1-mediated vascular endothelial growth factor expression. Mol Cell Biol 2001;21: 3995–4004.

(2) Bos R, van der Groep P, Greijer AE, Shvarts A, Meijer S, Pinedo HM, et al. Levels of hypoxia inducible factor-1alpha independently predict prognosis in patients with lymph node negative breast carcinoma. Cancer 2003;97: 1573–1581.

(3) Liu ZJ, Semenza GL, Zhang HF. Hypoxia-inducible factor 1 and breast cancer metastasis. J Zhejiang Univ Sci B 2015;16: 32–43.

(4) Sullivan R, Graham CH. Hypoxia-driven selection of the metastatic phenotype. Cancer Metastasis Rev 2007;26: 319–331.

(5) Semenza GL. Molecular mechanisms mediating metastasis of hypoxic breast cancer cells. Trends Mol Med 2012;18: 534–543.

(6) Flugel D, Gorlach A, Kietzmann T. GSK-3beta regulates cell growth, migration, and angiogenesis via Fbw7 and USP28-dependent degradation of HIF-1alpha. Blood 2012;119: 1292–1301.

(7) Luo W, Zhong J, Chang R, Hu H, Pandey A, Semenza GL. Hsp70 and CHIP selectively mediate ubiquitination and degradation of hypoxia-inducible factor (HIF)-1alpha but Not HIF-2alpha. J Biol Chem 2010;285: 3651–3663.

(8) Myllyharju J. Prolyl 4-hydroxylases, master regulators of the hypoxia response. Acta Physiol (Oxf) 2013;208: 148–165.

(9) Ivan M, Haberberger T, Gervasi DC, Michelson KS, Gunzler V, Kondo K, et al. Biochemical purification and pharmacological inhibition of a mammalian prolyl hydroxylase acting on hypoxia-inducible factor. Proc Natl Acad Sci U S A 2002;99: 13459–13464.

(10) Jaakkola P, Mole DR, Tian YM, Wilson MI, Gielbert J, Gaskell SJ, et al. Targeting of HIF-alpha to the von Hippel-Lindau ubiquitylation complex by O2-regulated prolyl hydroxylation. Science 2001;292: 468–472.

(11) Okumura F, Matsuzaki M, Nakatsukasa K, Kamura T. The role of Elongin BC-containing ubiquitin ligases. Front Oncol 2012;2.

(12) Soubeyran P, Kowanetz K, Szymkiewicz I, Langdon WY, Dikic I. Cbl-CIN85-endophilin complex mediates ligand-induced downregulation of EGF receptors. Nature 2002;416: 183–187.

(13) Kowanetz K, Husnjak K, Höller D, Kowanetz M, Soubeyran P, Hirsch D, et al. CIN85 associates with multiple effectors controlling intracellular trafficking of epidermal growth factor receptors. Mol Biol Cell 2004;15: 3155–3166.

(14) Minegishi Y, Shibagaki Y, Mizutani A, Fujita K, Tezuka T, Kinoshita M, et al. Adaptor protein complex of FRS2ß and CIN85/CD2AP provides a novel mechanism for ErbB2/HER2 protein downregulation. Cancer Sci 2013;104: 345–352.

(15) Petrelli A, Gilestro GF, Lanzardo S, Comoglio PM, Migone N, Giordano S. The endophilin-CIN85-Cbl complex mediates ligand-dependent downregulation of c-Met. Nature 2002;416: 187–190.

(16) Kim J, Kang D, Sun BK, Kim J-, Song JJ. TRAIL/MEKK4/p38/HSP27/Akt survival network is biphasically modulated by the Src/CIN85/c-Cbl complex. Cell Signal 2013;25: 372–379.

(17) Narita T, Amano F, Yoshizaki K, Nishimoto N, Nishimura T, Tajima T, et al. Assignment of SH3KBP1 to human chromosome band Xp22.1→p21.3 by in situ hybridization. Cytogenet Cell Genet 2001;93: 133–134.

(18) Havrylov S, Redowicz MJ, Buchman VL. Emerging roles of Ruk/CIN85 in vesicle-mediated transport, adhesion, migration and malignancy. Traffic 2010;11: 721–731.

(19) Nam JM, Onodera Y, Mazaki Y, Miyoshi H, Hashimoto S, Sabe H. CIN85, a Cbl-interacting protein, is a component of AMAP1-mediated breast cancer invasion machinery. EMBO J 2007;26: 647–656.

(20) Samoylenko A, Vynnytska-Myronovska B, Byts N, Kozlova N, Basaraba O, Pasichnyk G, et al. Increased levels of the HER1 adaptor protein Rukl/CIN85 contribute to breast cancer malignancy. Carcinogenesis 2012;33: 1976–1984.

(21) Wakasaki T, Masuda M, Niiro H, Jabbarzadeh-Tabrizi S, Noda K, Taniyama T, et al. A critical role of c-Cbl-interacting protein of 85 kDa in the development and progression of head and neck squamous cell carcinomas through the Ras-ERK pathway. Neoplasia 2010;12: 789–796.

(22) Cascio S, Finn OJ. Complex of MUC1, CIN85 and Cbl in colon cancer progression and metastasis. Cancers 2015;7: 342–352.

(23) Samoylenko A, Dimova EY, Kozlova N, Drobot L, Kietzmann T. The adaptor protein Ruk/CIN85 activates plasminogen activator inhibitor-1 (PAI-1) expression via hypoxia-inducible factor-1alpha. Thromb Haemost 2010;103: 901–909.

(24) Berra E, Benizri E, Ginouves A, Volmat V, Roux D, Pouyssegur J. HIF prolyl-hydroxylase 2 is the key oxygen sensor setting low steady-state levels of HIF-1alpha in normoxia. EMBO J 2003;22: 4082–4090.

(25) Kerppola TK. Design and implementation of bimolecular fluorescence complementation (BiFC) assays for the visualization of protein interactions in living cells. Nat Protoc 2006;1: 1278–1286.

(26) Kerppola TK. Bimolecular fluorescence complementation (BiFC) analysis as a probe of protein interactions in living cells. Annu Rev Biophys 2008;37: 465–487.

(27) Bruick RK, McKnight SL. A conserved family of prolyl-4-hydroxylases that modify HIF. Science 2001;294: 1337–1340.

(28) Appelhoff RJ, Tian YM, Raval RR, Turley H, Harris AL, Pugh CW, et al. Differential function of the prolyl hydroxylases PHD1, PHD2, and PHD3 in the regulation of hypoxia-inducible factor. J Biol Chem 2004;279: 38458–38465.

(29) Ma Y, Ye F, Xie X, Zhou C, Lu W. Significance of PTPRZ1 and CIN85 expression in cervical carcinoma. Arch Gynecol Obstet 2011;284: 699–704.

(30) Schroeder B, Weller SG, Chen J, Billadeau D, McNiven MA. A Dyn2-CIN85 complex mediates degradative traffic of the EGFR by regulation of late endosomal budding. EMBO J 2010;29: 3039–3053.

(31) Yakymovych I, Yakymovych M, Zang G, Mu Y, Bergh A, Landström M, et al. CIN85 modulates TGFB signaling by promoting the presentation of TGFB receptors on the cell surface. J Cell Biol 2015;210: 319–332.

(32) Cascio S, Farkas AM, Hughey RP, Finn OJ. Altered glycosylation of MUC1 influences its association with CIN85: The role of this novel complex in cancer cell invasion and migration. Oncotarget 2013;4: 1700–1711.

(33) Schmidt MHH, Chen B, Randazzo LM, Bögler O. SETA/CIN85/Ruk and its binding partner AIP1 associate with diverse cytoskeletal elements, including FAKs, and modulate cell adhesion. J Cell Sci 2003;116: 2845–2855.

(34) Hernandez-Valladares M, Kim T, Kannan B, Tung A, Aguda AH, Larsson M, et al. Structural characterization of a capping protein interaction motif defines a family of actin filament regulators. Nat Struct Mol Biol 2010;17: 497–503.

(35) Dunwoodie SL. The Role of Hypoxia in Development of the Mammalian Embryo. Dev Cell 2009;17: 755–773.

(36) Dales J-, Garcia S, Meunier-Carpentier S, Andrac-Meyer L, Haddad O, Lavaut M-, et al. Overexpression of hypoxia-inducible factor HIF-1a predicts early relapse in breast cancer: Retrospective study in a series of 745 patients. Int J Cancer 2005;116: 734–739.

(37) Zhong H, Agani F, Baccala AA, Laughner E, Rioseco-Camacho N, Isaacs WB, et al. Increased expression of hypoxia inducible factor-1alpha in rat and human prostate cancer. Cancer Res 1998;58: 5280–5284.

(38) Koivunen P, Tiainen P, Hyvarinen J, Williams KE, Sormunen R, Klaus SJ, et al. An endoplasmic reticulum transmembrane prolyl 4-hydroxylase is induced by hypoxia and acts on hypoxia-inducible factor alpha. J Biol Chem 2007;282: 30544–30552.

(39) Jokilehto T, Högel H, Heikkinen P, Rantanen K, Elenius K, Sundström J, et al. Retention of prolyl hydroxylase PHD2 in the cytoplasm prevents PHD2-induced anchorage-independent carcinoma cell growth. Exp Cell Res 2010;316: 1169–1178.

(40) Taylor MS. Characterization and comparative analysis of the EGLN gene family. Gene 2001;275: 125–132.

(41) Stolze IP, Mole DR, Ratcliffe PJ. Regulation of HIF: Prolyl hydroxylases. Novartia Found Symp 2006;272: 15–25.

(42) Colla S, Tagliaferri S, Morandi F, Lunghi P, Donofrio G, Martorana D, et al. The new tumor-suppressor gene inhibitor of growth family member 4 (ING4) regulates the production of proangiogenic molecules by myeloma cells and suppresses hypoxia-inducible factor-1 alpha (HIF-1alpha) activity: involvement in myeloma-induced angiogenesis. Blood 2007;110: 4464–4475.

(43) Song D, Li L-, Heaton-Johnson KJ, Arsenault PR, Master SR, Lee FS. Prolyl hydroxylase domain protein 2 (PHD2) binds a Pro-Xaa-Leu-Glu motif, linking it to the heat shock protein 90 pathway. J Biol Chem 2013;288: 9662–9674.

(44) Lee SH, Bae SC, Kim KW, Lee YM. RUNX3 inhibits hypoxia-inducible factor-1a protein stability by interacting with prolyl hydroxylases in gastric cancer cells. Oncogene 2014;33: 1458–1467.

(45) Baek JH, Mahon PC, Oh J, Kelly B, Krishnamachary B, Pearson M, et al. OS-9 interacts with hypoxia-inducible factor 1alpha and prolyl hydroxylases to promote oxygen-dependent degradation of HIF-1alpha. Mol Cell 2005;17: 503–512.

(46) Foxler DE, Bridge KS, James V, Webb TM, Mee M, Wong SC, et al. The LIMD1 protein bridges an association between the prolyl hydroxylases and VHL to repress HIF-1 activity. Nat Cell Biol 2012;14: 201–208.

(47) Zhang C-, Liu Q, Li M, Lin S-, Peng Y, Wu D, et al. RHOBTB3 promotes proteasomal degradation of HIFa through facilitating hydroxylation and suppresses the Warburg effect. Cell Res 2015;25: 1025–1042.

(48) Yu Y-, Chen C, Kong F-, Zhang W. Clinicopathological significance and potential drug target of RUNX3 in breast cancer. Drug Des Dev Ther 2014;8: 2423–2430.

(49) Ozer A, Bruick RK. Regulation of HIF by prolyl hydroxylases: Recruitment of the candidate tumor suppressor protein ING4. Cell Cycle 2005;4: 1153–1156.

(50) Sharp TV, Munoz F, Bourboulia D, Presneau N, Darai E, Wang H-, et al. LIM domains-containing protein 1 (LIMD1), a tumor suppressor encoded at chromosome 3p21.3, binds pRB and represses E2F-driven transcription. Proc Natl Acad Sci U S A 2004;101: 16531–16536.

(51) Pientka FK, Hu J, Schindler SG, Brix B, Thiel A, Jöhren O, et al. Oxygen sensing by the prolyl-4-hydroxylase PHD2 within the nuclear compartment and the influence of compartmentalisation on HIF-1 signalling. J Cell Sci 2012;125: 5168–5176.

(52) Zarrinpar A, Bhattacharyya RP, Lim WA. The structure and function of proline recognition domains. Sci STKE 2003;2003.

(53) Koch CA, Anderson D, Moran MF, Ellis C, Pawson T. SH2 and SH3 Domains: Elements that control interactions of cytoplasmic signaling proteins. Science 1991;252: 668–674.

(54) Kurakin AV, Wu S, Bredesen DE. Atypical Recognition Consensus of CIN85/SETA/Ruk SH3 Domains Revealed by Target-assisted Iterative Screening. J Biol Chem 2003;278: 34102–34109.

(55) Barth S, Edlich F, Berchner-Pfannschmidt U, Gneuss S, Jahreis G, Hasgall PA, et al. Hypoxia-inducible factor prolyl-4-hydroxylase PHD2 protein abundance depends on integral membrane anchoring of FKBP38. J Biol Chem 2009;284: 23046–23058.

(56) Barth S, Nesper J, Hasgall PA, Wirthner R, Nytko KJ, Edlich F, et al. The peptidyl prolyl cis/trans isomerase FKBP38 determines hypoxia-inducible transcription factor prolyl-4-hydroxylase PHD2 protein stability. Mol Cell Biol 2007;27: 3758–3768.

(57) Chowdhury R, Leung IK, Tian YM, Abboud MI, Ge W, Domene C, et al. Structural basis for oxygen degradation domain selectivity of the HIF prolyl hydroxylases. Nat Commun 2016;7: 12673.

(58) Choi KO, Lee T, Lee N, Kim JH, Yang EG, Yoon JM, et al. Inhibition of the catalytic activity of hypoxia-inducible factor-1alpha-prolyl-hydroxylase 2 by a MYND-type zinc finger. Mol Pharmacol 2005;68: 1803–1809.

(59) Kawata A, Iida J, Ikeda M, Sato Y, Mori H, Kansaku A, et al. CIN85 is localized at synapses and forms a complex with S-SCAM via dendrin. J Biochem 2006;139: 931–939.

(60) Hassinen A, Rivinoja A, Kauppila A, Kellokumpu S. Golgi N-glycosyltransferases form both homo- and heterodimeric enzyme complexes in live cells. J Biol Chem 2010;285: 17771–17777.

(61) Borthwick EB, Korobko IV, Luke C, Drel VR, Fedyshyn YY, Ninkina N, et al. Multiple domains of Ruk/CIN85/SETA/CD2BP3 are involved in interaction with p85a regulatory subunit of PI 3-kinase. J Mol Biol 2004;343: 1135–1146.

(62) Gout I, Middleton G, Adu J, Ninkina NN, Drobot LB, Filonenko V, et al. Negative regulation of PI 3-kinase by Ruk, a novel adaptor protein. EMBO J 2000;19: 4015–4025.

(63) Scharf JG, Unterman TG, Kietzmann T. Oxygen-dependent modulation of insulin-like growth factor binding protein biosynthesis in primary cultures of rat hepatocytes. Endocrinology 2005;146: 5433–5443.

(64) Stewart SA, Dykxhoorn DM, Palliser D, Mizuno H, Yu EY, An DS, et al. Lentivirus-delivered stable gene silencing by RNAi in primary cells. RNA 2003;9: 493–501.

(65) Sanjana NE, Shalem O, Zhang F. Improved vectors and genome-wide libraries for CRISPR screening. Nat Methods 2014;11: 783–784.

(66) Shalem O, Sanjana NE, Hartenian E, Shi X, Scott DA, Mikkelsen TS, et al. Genome-scale CRISPR-Cas9 knockout screening in human cells. Science 2014;343: 84–87.

(67) Kozlova N, Samoylenko A, Drobot L, Kietzmann T. Urokinase is a negative modulator of Egf-dependent proliferation and motility in the two breast cancer cell lines MCF-7 and MDA-MB-231. Mol Carcinog 2016;55: 170–181.

(68) Mayevska O, Shuvayeva H, Igumentseva N, Havrylov S, Basaraba O, Bobak Y, et al. Expression of adaptor protein Ruk/CIN85 isoforms in cell lines of various tissue origins and human melanoma. Exp Oncol 2006;28: 275–281.

(69) Maruyama T, Dougan SK, Truttmann MC, Bilate AM, Ingram JR, Ploegh HL. Increasing the efficiency of precise genome editing with CRISPR-Cas9 by inhibition of nonhomologous end joining. Nat Biotechnol 2015;33: 538–542.

(70) Chu VT, Weber T, Wefers B, Wurst W, Sander S, Rajewsky K, et al. Increasing the efficiency of homology-directed repair for CRISPR-Cas9-induced precise gene editing in mammalian cells. Nat Biotechnol 2015;33: 543–548.

(71) Zhu X, Xu Y, Yu S, Lu L, Ding M, Cheng J, et al. An efficient genotyping method for genome-modified animals and human cells generated with CRISPR/Cas9 system. Sci Rep 2014;4.

(72) Schmittgen TD, Livak KJ. Analyzing real-time PCR data by the comparative C(T) method. Nat Protoc 2008;3: 1101–1108.

(73) Tan SC, Carr CA, Yeoh KK, J. Schofield C, Davies KE, Clarke K. Identification of valid housekeeping genes for quantitative RT-PCR analysis of cardiosphere-derived cells preconditioned under hypoxia or with prolyl-4-hydroxylase inhibitors. Mol Biol Rep 2012;39: 4857–4867.

(74) Konzack A, Jakupovic M, Kubaichuk K, Gorlach A, Dombrowski F, Miinalainen I, et al. Mitochondrial Dysfunction Due to Lack of Manganese Superoxide Dismutase Promotes Hepatocarcinogenesis. Antioxid Redox Signal 2015;23: 1059–1075.

(75) Hirsila M, Koivunen P, Gunzler V, Kivirikko KI, Myllyharju J. Characterization of the human prolyl 4-hydroxylases that modify the hypoxia-inducible factor. J Biol Chem 2003; 278: 30772–30780.

(76) Tomayko MM, Reynolds CP. Determination of subcutaneous tumor size in athymic (nude) mice. Cancer Chemother Pharmacol 1989;24: 148–154.

